# The Archaellum of *Methanospirillum hungatei* is Electrically Conductive

**DOI:** 10.1101/458356

**Authors:** David J.F. Walker, Eric Martz, Dawn E. Holmes, Zimu Zhou, Stephen S. Nonnenmann, Derek R. Lovley

## Abstract

Here we report that the archaellum of *Methanospirillum hungatei* is electrically conductive. Our analysis of the previously published archaellum structure suggests that a core of tightly packed phenylalanines is one likely route for electron conductance. This is the first demonstration that electrically conductive protein filaments (e-PFs) have evolved in Archaea and is the first e-PF for which a structure is known, facilitating mechanistic evaluation of long-range electron transport in e-PFs.

---

Electrically conductive pili (e-pili) expressed by microbes in the domain Bacteria play an important role in extracellular electron exchange between cells and their extracellular environment ^1,2^ e-Pili are found in diverse bacteria ^1,3^ but have been studied most extensively in *Geobacter sulfurreducens*, and related *Geobacter* species, in which e-pili are essential for long-range electron transport to Fe(III) oxide minerals, interspecies electron transfer, and electron conduction through biofilms ^1^. e-Pili enable unprecedented long-range (μm) electron conduction along the length of a protein filament, which not only has important biological implications, but also suggests diverse applications for these ‘protein nanowires’ as a sustainably produced electronic material ^1,4–6^. There is substantial debate over the potential mechanisms of long-range electron transport in e-pili ^1,5,6^, which is unresolved in large part because of the lack of a definitive e-pili structure. Although it has been possible to determine the structure of some pili with cryo-electron microscopy (cryo-EM)^7^, to date, the thin diameter (3 nm) and high flexibility of *G. sulfurreducens* e-pili have thwarted structural determination with the cryo-EM approach.

The finding that e-pili have independently evolved multiple times in Bacteria ^3^, raised the question of whether conductive protein filaments have ever evolved in Archaea. Diverse Archaea exchange electrons with their extracellular environment, reducing extracellular electron acceptors or engaging in direct interspecies electron transfer (DIET) with bacteria^28^. The alpha-helix filament structure of archaella resembles that of type IV pili ^7,9,10^, suggesting at least a remote possibility of producing an electrically conductive archaellum (e-archaellum) with properties similar to e-pili.

We chose the methanogen *Methanospirillum hungatei* for the initial search for an e-archaellum because *M. hungatei* is capable of reducing extracellular electron acceptors ^11^; archaellum expression is readily induced in *M. hungatei*^12^; and a cryo-EM (3.4 Å) structure of the archaellum is available ^10^. *M. hungatei* cells were drop-cast on highly ordered pyrolytic graphite (HOPG) and examined with conductive atomic force microscopy, in which a conductive tip serves as a translatable top electrode. Cells with a polar archaellum with the expected height of 10 nm ^10^ were readily detected with topographic imaging in contact mode (Fig. 1a,b,d). Conductive imaging demonstrated that the archaellum was electrically conductive (Fig. 1c, d, e). Point-mode current-voltage (I-V) spectroscopy revealed a linear-like response with currents that were higher than at the same voltage with *G. sulfurreducens* e-pili prepared in the same manner (Fig. 1e). The pili of *G. sulfurreducens* strain Aro-5, which produces pili specifically designed for low conductivity ^13,14^, exhibited very low currents (Figure 1e).

**Fig. 1.**
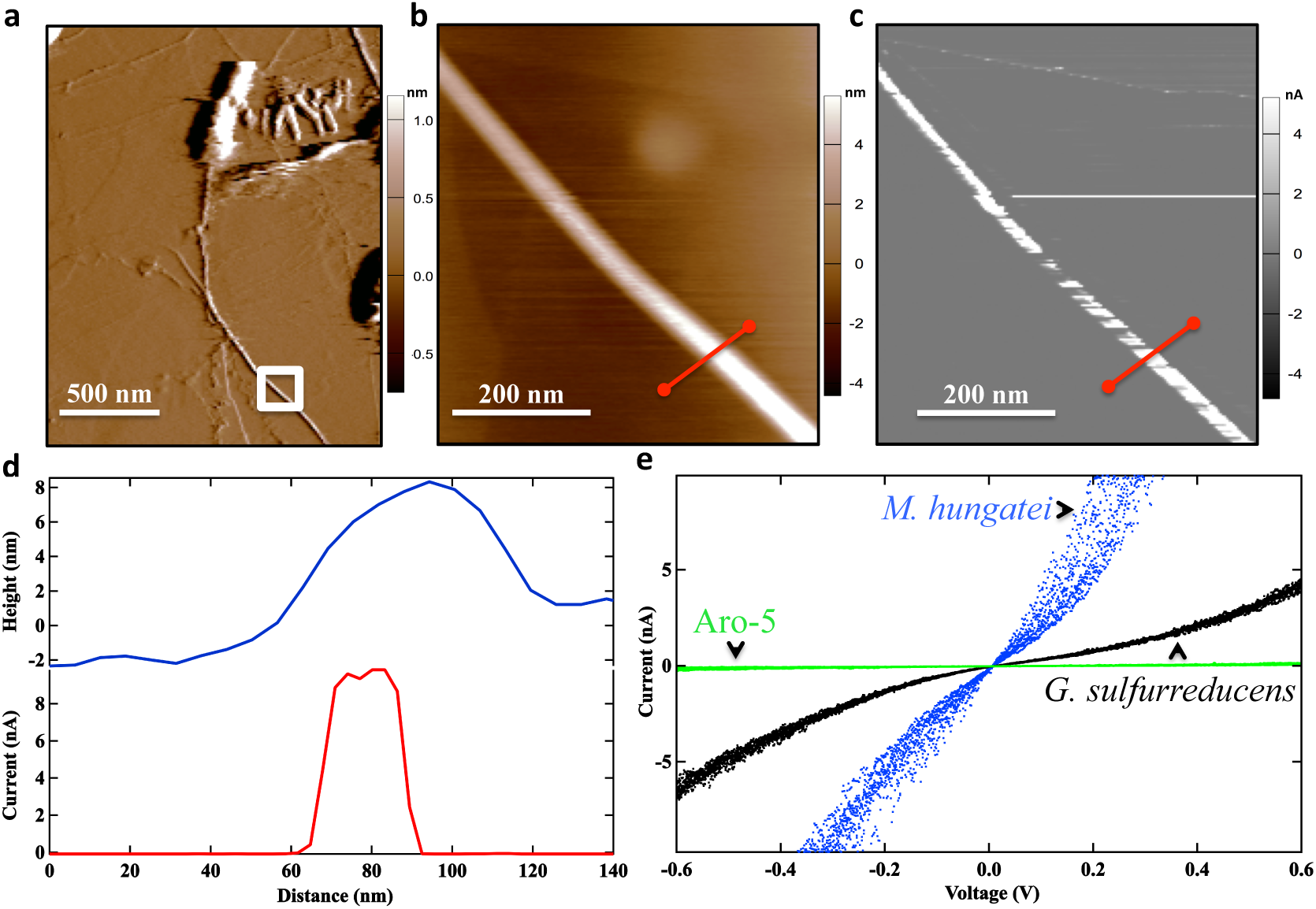
Electrical conductivity of the *Methanospirillum hungatei* archaellum determined with atomic force microscopy. **(a)** Contact topographic imaging of *M. hungatei* showing the polar archaellum protruding from the cell. The image was collected in deflection mode. The white box designates the region chosen for additional analysis. **(b)** Higher resolution topographic image of the archaellum from the region designated in (a). The red line indicates the slice taken for the topographic height and current cross-sectional line profile analysis. **(c)** Local current image of the individual archaellum with an applied bias of 300 mV. **(d)** Topographic height and current response from the cross sectional slice designated in (b). **(e)** Point-mode current response (I-V) spectroscopy of the individual archaellum (blue). Similar I-V analysis of the wild-type e-pili of *G. sulfurreducens* (black) and the poorly conductive pili of *G. sulfurreducens* strain Aro-5 (green) is shown for comparison. The *M. hungatei* archaellum conductivity measurement shown is representative of multiple measurements on multiple archaella (See Supplementary Figure 1 for additional examples).

The cellular electrical contacts for extracellular electron exchange in Archaea have yet to be elucidated. The discovery of an e-archaellum expands the possibilities to be considered. For example, e-archaella offer a potential mechanism for electrons derived from external sources to transverse the S-layer to interact with intracellular electron carriers and vice versa. Unfortunately, the current lack of genetic tools for manipulating *M. hungatei* prevents further evaluation of its physiological role with approaches, such as expressing a synthetic archaellin that yields a poorly conductive filament, that have been important for demonstrating the role of e-pili in *Geobacter* species ^1,13^

The cryo-EM structure of the *M. hungatei* e-archaellum ^10^ provides the first opportunity to directly evaluate possible routes for long-range electron transport along a biologically produced protein filament. Multiple lines of experimental evidence suggest that closely packed aromatic amino acids confer conductivity to *G. sulfurreducens* e-pili ^1,5^ Analysis of the distribution of aromatic amino acids in the *M. hungatei* e-archaellum revealed aromatic side chains organized in three distinct groups: a core, a middle sleeve, and an outer sleeve (Fig. 2a). The aromatic rings in the middle and outer sleeves appear too widely spaced to support conductivity (Fig. 2c,d). However, phenylalanine rings in the core are packed almost as close as is physically possible ^15^. The distances between ring centers of Phe1 and Phe13 are 4.5 and 5.1 Å (Fig. 2e), which is conceivably close enough for π-π interactions, similar to the π-π interactions proposed to be involved in electron conduction along *G. sulfurreducens* e-pili ^16–18^.

**Fig. 2.**
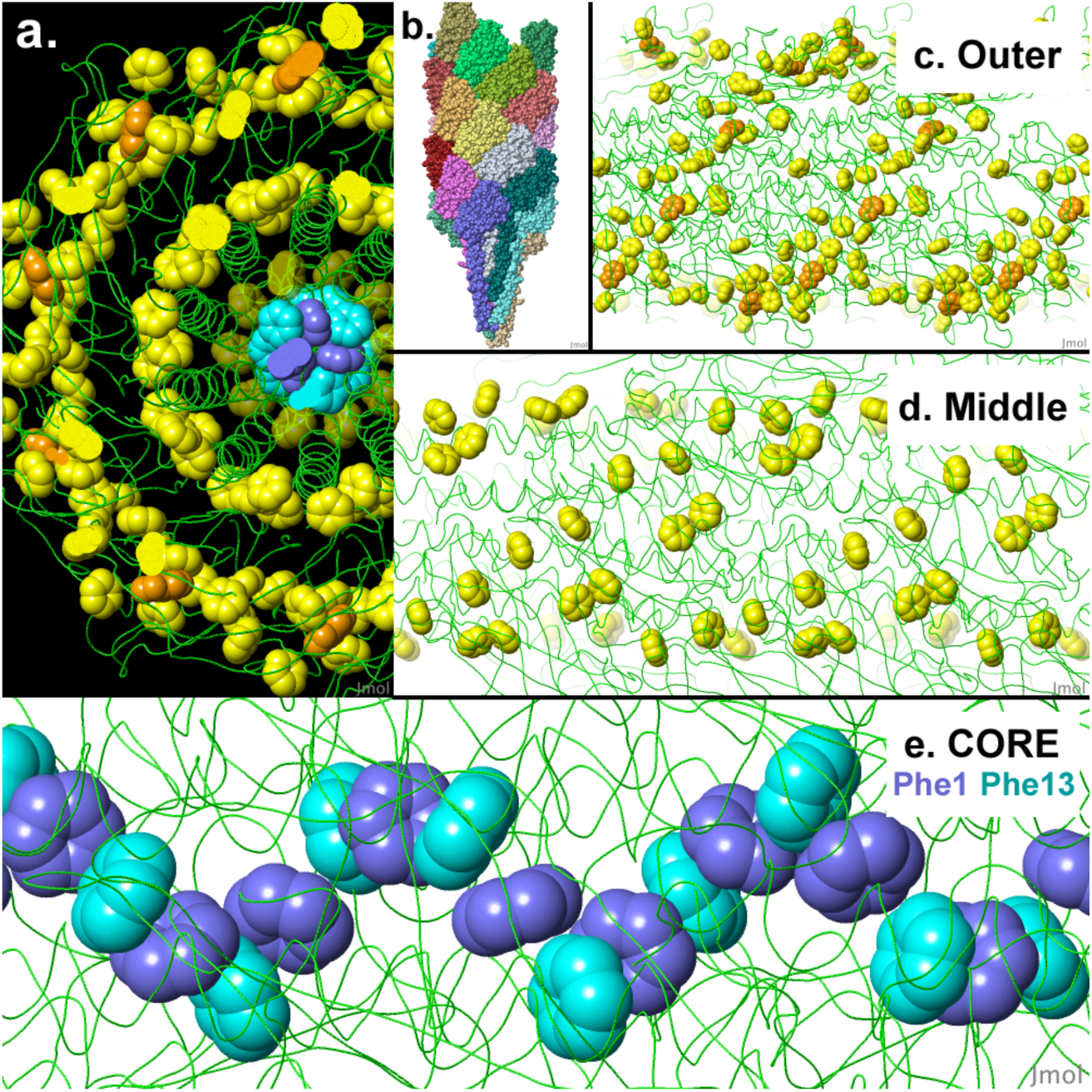
Tightly-packed aromatic rings form the core of the previously determined structure (PDB code 5tfy ^10^) of the *M. hungatei* archaellum. **(a)** In cross section, aromatic rings form three well-separated groups: a core (Phe1 blue, Phe13 cyan, Phe20 dim yellow), a middle sleeve, and an outer sleeve (Phe and Tyr yellow; Trp orange). **(b)** The entire model is an assembly of 26 protein chains (all atoms shown, each chain a distinct color, axis vertical). **(c)** and **(d)**, Side views (axis horizontal) of outer and middle sleeves of aromatics (rear half of model hidden). **(e)** Tightly packed core of alternating Phe1 and Phe13 rings. Ring center distances are 4.5 and 5.1 Å. Images and measurements made with Jmol.Org. See Supplementary Figure 2 for animations.

We suggest that this phenylalanine core is at least one of the features contributing to the e-archaellum conductivity, consistent with recent experimental evidence that has suggested that phenylalanines within the hydrophobic core of an amino acid α-helical structure can facilitate long-range electron transport ^19,20^. Genetic manipulations to alter the position of phenylalanines within the *M. hungatei* archaellum, analogous to the approach that has been used to evaluate the role of aromatic amino acid stacking in *G. sulfurreducens* e-pili ^13,14,17,18^, would provide a further test of this hypothesis. The added benefit of such studies with the *M. hungatei* e-archaellum is that it will be possible to directly examine structural modifications to electron conductance pathways with cryo-EM. In the absence of genetic tools for*M. hungatei*, it will be necessary to heterologously express the gene for the *M. hungatei* archaellin in a genetically tractable archael host, similar to the expression of heterologous e-pili in *G. sulfurreducens* ^3^, or to identify a similar e-archaellum in a genetically tractable archaeon.

Microbially produced protein nanowires show substantial promise as a sustainable “green” electronic material with possibilities for functionalization and biocompatibility not available with other nanowire materials ^1,4–6^. e-Archaella offer a unique opportunity directly examine how synthetic designs to tune conductivity and/or add functionality influences protein nanowire structure, enabling a less empirical approach to the design of protein nanowire electronics.

The discovery of e-archaella indicates that a search for electrically conductive protein filaments in other Archaea as well as the Eukarya is warranted. The high energetic cost for the biosynthesis of the abundant aromatic amino acids necessary to produce conductive filaments suggests positive selection for conductivity. For microbes like *Geobacter*, the benefit in promoting extracellular electron exchange is clear. The possibility that other physiological functions, such electrical signaling between cells, may have provided an evolutionary advantage in *M. hungatei* and other organisms should be explored.

## Methods

*M. hungatei* was grown as previously described ^12^ in low phosphate medium to induce archaellum expression. An aliquot (100 μl) of the culture was drop-cast onto highly oriented pyrolytic graphite (HOPG). Cells were allowed to attach to the HOPG for 10 min and then the liquid was removed with a pipette tip. The surface was washed twice with 100 μl of deionized water, the surface was blotted dry at the edge with a Kimwipe, and placed in a desiccator overnight. Samples were equilibrated with atmospheric humidity for at least two hours. Conductive atomic force microscopy was performed using an Oxford Instruments/Asylum Research Cypher ES atomic force microscope. All topographic and current imaging was performed with a Pt/Ir-coated Arrow-ContPT tip with a 0.2 N/m force constant (NanoWorld AG, Neuchâtel, Switzerland). Topographic imaging required a set point of 0.002V. The conductive tip was attached to an ORCA™ dual-gain transimpedance amplifier and held at ground to serve as a translatable top electrode. A 300 mV bias was applied to the HOPG and the locally detected current response of the archaellum was identified. Point-mode current-voltage (I-V) spectroscopy was performed by applying the conducting AFM tip to the top of the archaellum (0.002V) and performing a voltage sweep at a frequency of 0.99 Hz.

## Acknowledgments

We thank Trevor Woodard for growing the *Methanospirillum hungatei cultures*. This research was supported by Office of Naval Research grant N00014-16-1-2526.

## Author contributions

D.J.F.W., D.E.H., and D.R.L. conceived the project, D.J.F.W. performed the atomic force microscopy measurements with assistance and guidance from Z.Z. and S.S.N. E.M. performed the analysis of the structural model. D.J.F.W. and D.R.L. wrote the initial draft of the manuscript with comments revisions contributed by all of the authors.

